# The use of plant extracts and bacteriophages as an alternative therapy approach in combating bacterial infections: the study of lytic phages and *Stevia rebaudiana*

**DOI:** 10.1101/2023.06.27.546765

**Authors:** Xymena Stachurska, Małgorzata Mizielińska, Magdalena Ordon, Paweł Nawrotek

## Abstract

**Introduction:** In the light of the problem of antibiotic-resistance, the use of alternative combined therapies in combating bacterial-related issues has gained popularity. Therefore, using up-to-date laboratory techniques, possible interactions of lytic bacteriophages (MS2, T4 and Phi6), acetone and methanol *S. rebaudiana* extracts (SRa and SRm) in the bacterial environment have been examined.

**Material and Methods:** Using a microdilution method, phages-extracts coincubation assay, static interactions synographies and dynamic growth profile experiments in a bioreactor, it was found that the interactions in a static environment differ from interactions in a dynamic environment.

**Results:** Dynamic conditions alter the influence of extracts in a concentration-dependent manner. The effects of the SRa and SRm extracts on bacterial growth in a dynamic environment depend on the species of the phage and bacterial host. The greatest differences were observed for *E. coli* strains and their phages, whereas *P. syringae* and Phi6 phage reacted very similar to both extracts. The differences also emerged for single extracts within *E. coli* strains and their phages.

**Conclusions:** Every extract type should be tested on a case-by-case basis and experiments outcomes should not be generalized before gathering data. Moreover, many varied experiments should be performed, especially when examining such multifactorial mixtures. Tested mixtures could be potentially used in multi-drug resistant bacterial infections treatments.

## Introduction

Antibiotic resistance is a severe threat to the worldwide health, with substantial repercussions for the health of humans and animals, as well as food safety. Antibiotics are used excessively in human and animal treatment. Multiple antibiotic-resistant bacteria are a challenge in treating bacterial infections and their transmission from animal to human and from human to human that puts the health sector at risk. In the light of the emergence of resistant strains of microbes that cannot be effectively eradicated by antibiotics, exploring alternative therapy methods could be of great importance. Furthermore, antimicrobial resistance is still increasing worldwide (despite new treatments being employed) with a simultaneous decrease in the discovery rate of novel antibiotics (3, 30). Herbs and extracted compounds of plant origin are defined as phytogenics, phytobiotics or botanical natural additives. Numerous studies on these kinds of active compounds have shown their beneficial effects, such as antibacterial, antiviral or antioxidant (2, 3, 6, 9, 12). These substances have many advantages over commonly used antibiotics -they may be a replacement in the treatment of resistant bacteria infections.

*Stevia rebaudiana* is a herbaceous perennial plant of the genus *Stevia* Cav. and *Asteraceae* family. The leaves of stevia produce diterpene glycosides (stevioside and rebaudiosides). They are a non-nutritive, non-toxic, high-potency sweeteners and may serve as substitutes to sucrose as well as other synthetic sweeteners (44). The sweet compounds of stevia do not chemically break down as they pass through the human digestive system. Therefore, stevia is considered safe for those, whose sugar level in blood must be controlled (36, 13). Moreover, stevia can be grown very easily like any other vegetable crop, which makes it an easy-to-obtain plant (13) and it is also known to be free from insects. Therefore, insecticides are not required in its cultivation, which is of great help in producing organic stevia (44). In addition to its sweetening properties, it has medicinal values and uses. Literature points out other medical applications like antihypertensive, antihyperglycaemic, antitumour, antiinflammatory, antidiarrhoeal, diuretic, hepatoprotective and immunomodulatory effects (44, 9). Moreover, stevia leaves and callus have strong antioxidant activity and may be rich source of antioxidants (38).

The particular importance of stevia in the light of this research, is attached to its antimicrobial activity, especially against certain bacteria. However, it is still unclear whether stevia extracts influence bacteria positively or negatively (31, 20, 29). Furthermore, it has been shown that the antimicrobial potential of stevia depends on the type of used extract (different extraction methods) (33). The importance of any knowledge on the antibacterial action of stevia appears even greater in the light of recent studies, showing that combinations of naturally antimicrobial plant extracts with other active agents of similar effectiveness, may be more effective together than used as individual therapies (26).

Bacteriophages (phages) could serve as alternative antimicrobial agents that can potentially be combined with plant extracts to create more effective preparations. Phages are viruses that infect their specific bacterial hosts and can successfully supplement, or in some cases, even replace traditional methods of treatment and prevention of bacterial diseases. Especially when taking into consideration those diseases that are caused by antibiotic-resistant bacterial strains (1). Phages have been known for over a hundred years and today are regaining popularity once more. Phage virion consists of the molecules characterised by their biochemical complexity. Together with their diversified size, hydrophobicity and electric charge, bacteriophages interact with other materials and factors in various ways, including stimulating or inhibiting effects. An example would be the alteration of a phage’s lytic activity as a result of synergistic or antagonistic influence (45, 34). Moreover, their easy handling and ability to be cultivated in standard laboratory media are significant advantages that make bacteriophages easy to “produce”. Additionally, it has been proposed that due to these benefits, phages can be employed as eukaryotic cell viruses surrogates (28, 10, 41).

Therefore, in the present study, interactions between lytic bacteriophages and two different types of *Stevia rebaudiana* plant extract (methanol and acetone) were investigated for their possible relationships in a bacterial environment, including any synergistic or antagonistic effects. To the best of our knowledge, this is the first work describing such phenomena.

## Material and Methods

### Microorganisms

Bacteriophages and their corresponding bacterial hosts were purchased from a German Collection of Microorganisms and Cell Cultures GmbH (Deutsche Sammlung von Mikroorganismen und Zellkulturen; DSMZ). *Pseudomonas syringae* (DSM 21482) with phage Phi6 (DSM 21518), *Escherichia coli* (DSM 5695) with phage MS2 (DSM 13767) and *Escherichia coli* (DSM 613) with phage T4 (DSM 4505) were used as model microorganisms based on their different features presented below (Table 1).

**Table 1.**
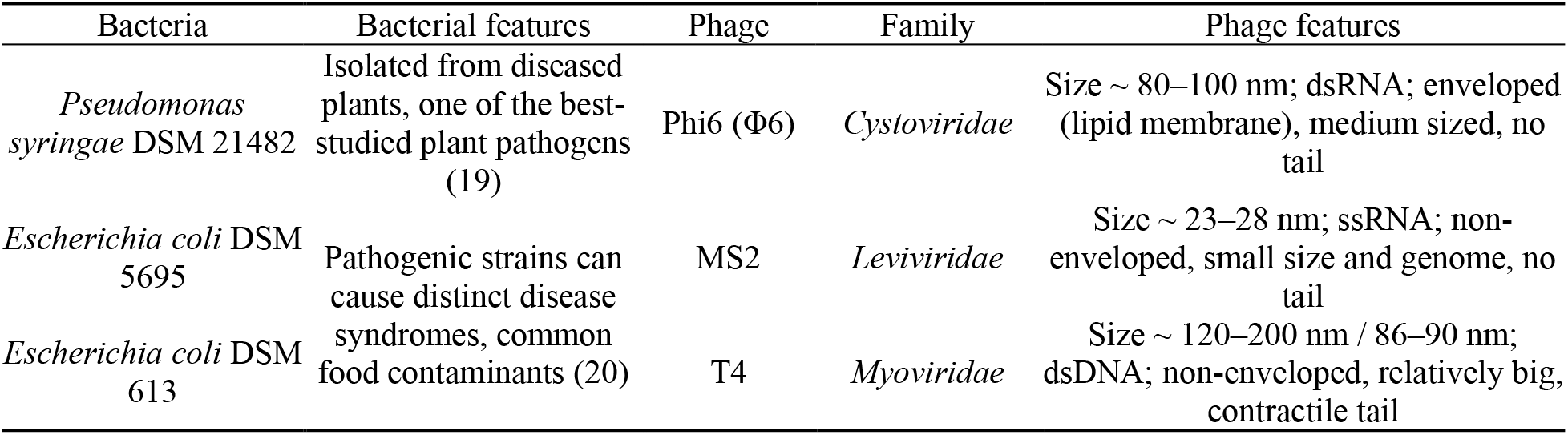
Characteristics of the tested microorganisms

Bacterium revival and phage propagation was conducted as described earlier (35). Briefly, the bacteria were revived from storage at –20 °C in trypticase soy broth (BioMaxima, Lublin, Poland) with 10 % (vol/vol) glycerol on Luria-Bertani (LB agar) (BioMaxima, Lublin, Poland) by streaking glycerol stocks onto agar plates. The plates were then incubated for 24 h at 37 °C – *E. coli* strains and for 48 h at 28 °C – *P. syringae*. Phages were amplified by inoculating 50 mL of LB with colonies from bacterial stock plates and incubated (as described above) with shaking (120 rpm) in an orbital rotating shaker (Shaker-Incubator ES-20, BioSan, Józefów, Poland) to reach optical density of 0.2 (OD_600nm_ = 0.2). OD was measured using an Infinite 200 PRO NanoQuant microplate reader (Tecan, Männedorf, Switzerland). At this point the phages were added and samples were further incubated until lysis occurred. For MS2 and T4 lysate purification, samples were supplemented with chloroform (10 %, vol/vol), vortexed for 5 min and then centrifuged (Eppendorf Centrifuge model 5810 R, Hamburg, Germany) at 5000 rpm for 25 min at 4 °C. The supernatant (phage lysate) was collected immediately and stored at 4 °C for further use. For Phi6 lysate purification the sample was centrifuged at the first stage (5000 rpm, 15 min, 4 °C) and then filter sterilized (PES filter, 0.22 µm). The lysate was collected and stored at 4 °C for further use. Phages activity and titres were tested by a double overlay agar plaque assay (18).

### The preparation of acetone and methanol extracts of *Stevia rebaudiana*

Preparation of *S. rebaudiana* extracts was carried out as described in our previous work (24) with some modifications. 50 g of dried herb leaves (Flos, Mokrsko, Poland) were placed in glass bottles. The first bottle was intended for methanol extraction, therefore 100 mL of the 70 % aqueous methanol (MeOH) was introduced into the first bottle containing dry herb leaves. Second bottle intended for acetone extraction was supplemented with 100 mL of the 70 % aqueous acetone. Next, the samples were placed in a shaker (Ika, Staufen im Breisgau, Germany) and extracted for 2 h at 70 °C with 150 rpm. The crude extracts were filtered through a Büchner funnel equipped with a cellulose filter. The extracts were then concentrated by evaporation at 50 °C to obtain 10 mL of each aqueous solution. After the evaporation of methanol and acetone, the samples were diluted with 20 mL of water and sterilized by filtration (PES, 0.22 µm). The samples were then used for further experiments. At this stage the *Stevia rebaudiana* water solution of active compounds obtained by methanol extraction was marked as “SRm” and the *Stevia rebaudiana* water solution of active substances obtained by acetone extraction was marked as “SRa”. Additionally, the dry mass of each extract was determined via moisture analyser (Radwag, Puszczykowo, Poland). For the next experiments, different stock solutions (two-fold serial dilutions) of the extracts were diluted in sterile, deionized water and kept at –20 °C until further analysis.

### The influence of *S. rebaudiana* extracts on bacterial cells

In order to test the influence of *S. rebaudiana* acetone (SRa) and methanol (SRm) extracts on bacterial cells, a 96-well microplate dilution experiment was performed. For the microplate method, modified MIC determination was used (42). The colonies from stock bacterial plates were used to inoculate falcon tubes with 30 mL of LB and incubated (37 °C for *E. coli* strains and 28 °C for *P. syringae*) with shaking (120 rpm) in an orbital rotating shaker till the cultures reached OD_600nm_ = 0.2. 96-well polystyrene flat-bottomed plates were supplemented with two-fold serial dilutions of SRa and SRm (50 µL; 50 % – 0.003 %) followed by the addition of bacterial suspensions (50 µL). Sterile deionized water added in place of extracts was used as the positive control (bacteria growth control). Plates were then placed in incubators for 24 h at temperatures appropriate for the tested bacteria. Optical density values were measured using an Infinite 200 PRO NanoQuant microplate reader. The experiment was conducted in triplicate.

### Coincubation studies of phages and *S. rebaudiana* extracts

For direct phage-extract interactions testing, a coincubation experiment was performed (34). The test was carried out in order to reveal the influence of extracts on bacteriophages plaque forming ability and phage titers. Briefly, 12-well flat bottom polystyrene plates were initially supplemented with 1 mL of phage lysates (Phi6 and T4 – 10^8^ PFU/mL, MS2 – 10^9^ PFU/mL). Then 1 mL of SRa and SRm were added to reach the final concentrations of 50 % – 0.049 % (two-fold serial dilutions). Extract-free deionized water with phage lysate was used as a positive control (phage control). The plates were then incubated at room temperature (21 °C) for 24 h without light access. Later, samples were titered in TM buffer (50 mM Tris-HCl, 10 mM MgSO_4_ at pH 7.5) by spotting 3 µL of each 10-fold dilution onto an LB plate already coated with a top agar layer (7 %) mixed with overnight bacterial culture (double-layer agar technique). The experiment was conducted in triplicate.

### The interaction stoichiometries of phages and *S. rebaudiana* extracts

Phage-extract synographies were performed as described elsewhere (14) with minor modifications. For test cultures preparation, 5 mL of the overnight cultures was diluted in LB, in order to achieve OD_600nm_ = 1 (approx. 1 × 10^9^ CFU/mL) and 100 µL of the suspensions were transferred into each well of the 96-well flat-bottomed plates. The plates were previously inoculated with varying concentration range of phages (50 µL, final 10^2^–10^8^ PFU/mL) and extracts (50 µL, final 25 % – 0.049 %), creating a checkerboard assay. Plates were incubated for 18 h in 37 °C for *E. coli*, and in 28 °C for *P. syringae*. Bacterial turbidity was determined by measuring OD_600nm_ values, using an Infinite 200 PRO NanoQuant microplate reader. Afterwards, resazurin assay was carried out for bacteria metabolic activity testing. The dye was added to the samples in wells (final 1 mg/mL) and plates were further incubated (*E. coli* – 3 h, *P. syringae* – 4.5 h) without light access. Then, fluorescence measurements were performed using a fluorescent plate reader (Synergy HTX, BioTek, Winooski, VT, USA) at 540 nm excitation and 590 nm emission. The experiment was conducted in triplicate.

### Phage infection and lysis profile experiments

Based on results from the static synograms experiment (no mixing conditions), we chose combinations in which interesting phenomena were detected (increased bacterium activity in resazurin assay with simultaneous OD measures showing a reduction in bacterial biomass), in order to analyse the influence of combined treatment of phage-extracts on bacterial hosts growth rate in real time in a dynamic environment (mixing conditions). Proper controls for results comparison (phage + extract maximal dose, extract maximal dose, bacterial growth control) were also applied (Table 2).

**Table 2.**
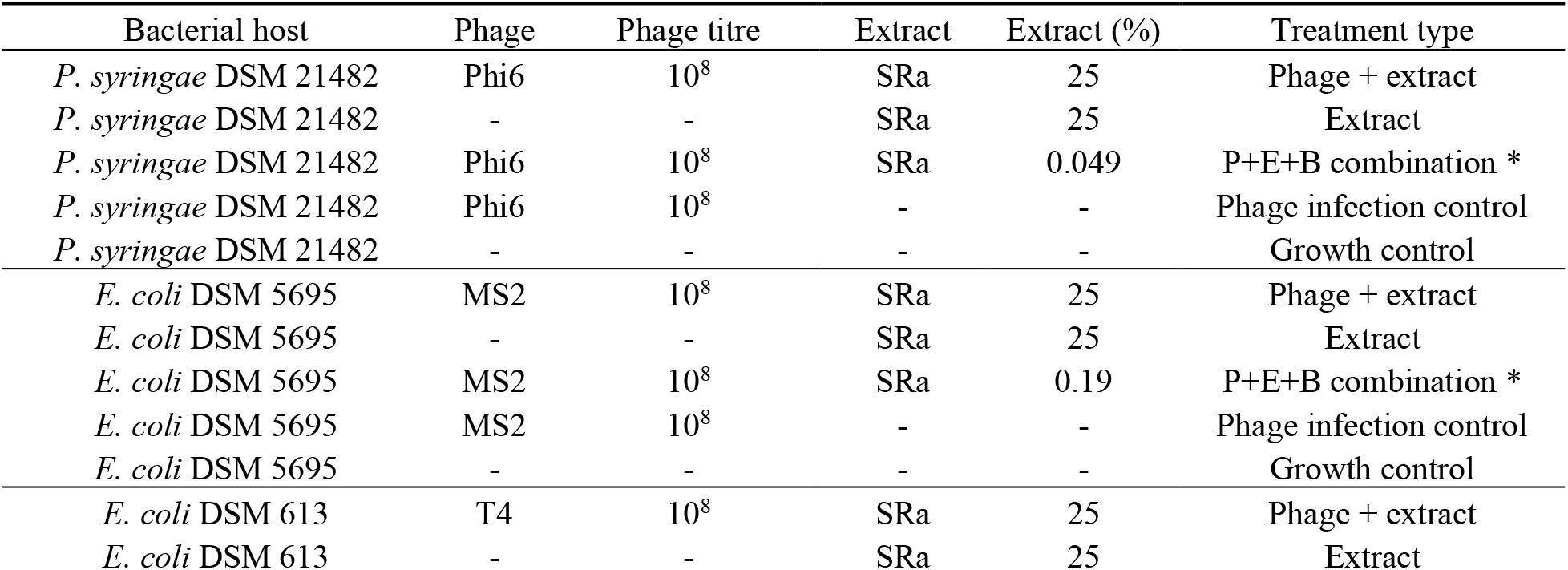
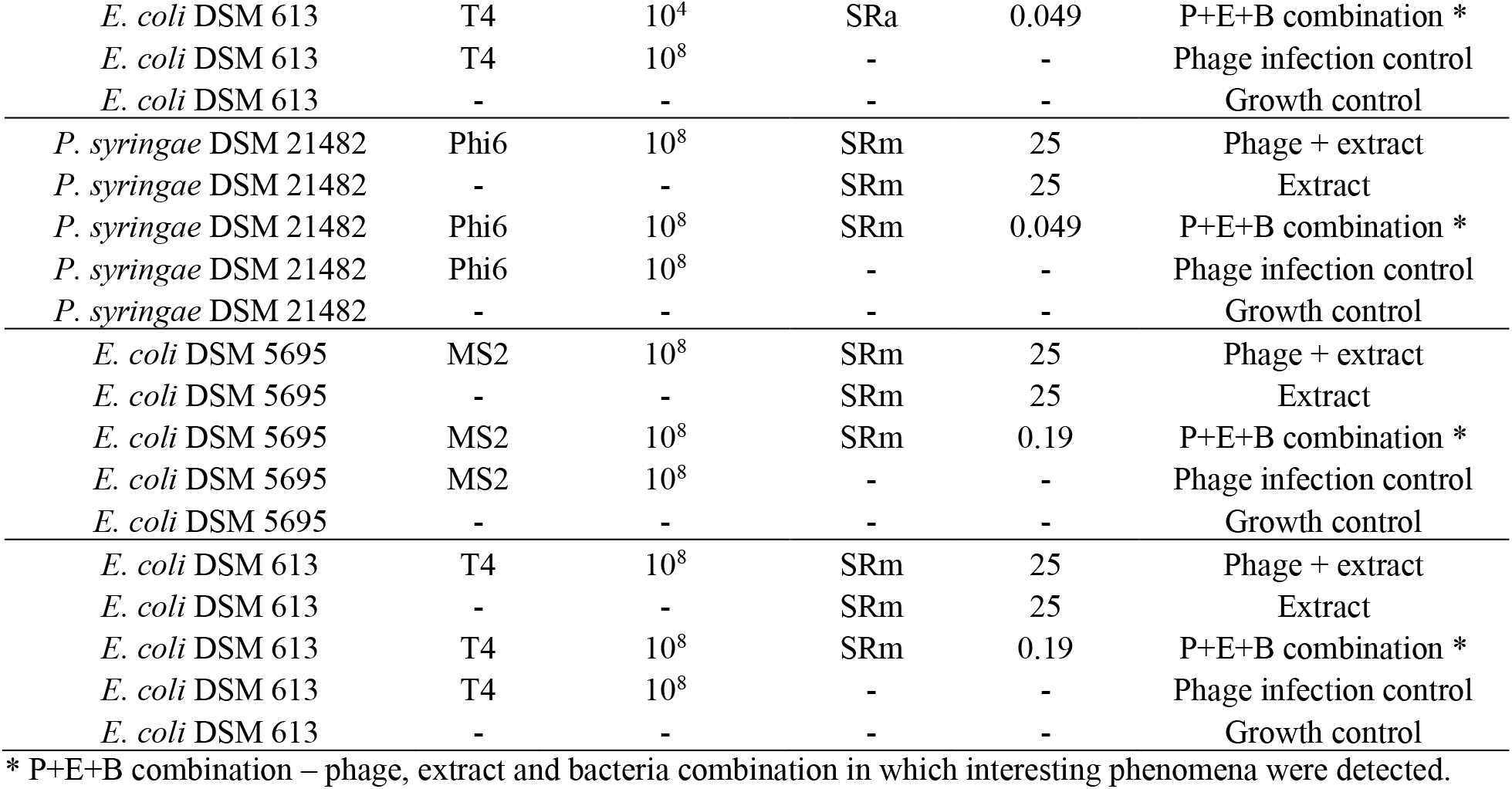
Chosen treatment combinations for real time assessment of bacterial growth. SRa – *S. rebaudiana* acetone extract; SRm – *S. rebaudiana* methanol extract

In order to keep the experimental assumptions of the synograms test, overnight bacterial host cultures were diluted in LB, in order to achieve OD_600nm_ = 1 and 5 mL of the suspensions were then transferred to 50 mL Falcon tubes. Afterwards phage lysates were added (2.5 mL, final titre 10^8^ PFU/mL) along with 2.5 mL of SRa or SRm extracts concentrations, in order to obtain the chosen final concentrations (e.g. 50 % extract was added to obtain a final concentration of 12.5 %). For extracts only treatments, instead of phage, 2.5 mL of sterile deionized water was added. For host bacterium growth control, 5 mL of sterile deionized water was added to 5 mL of the bacterial suspension. Samples were then incubated (*E. coli* – 37 °C, *P. syringae* – 28 °C) for 16 h, 150 rpm and real time OD_850nm_ values were measured using BioSan bioreactors (BS-010160-A04, BioSan, Riga, Latvia).

### Statistical analysis

One-way (plant extracts antimicrobial studies) and two-way (phage-extract coincubation assay) ANOVA was used to statistically analyse the results, along with Dunnett’s multiple comparisons test. Differences were considered significant at p ≤0.05. All statistical analyses were carried out using GraphPad Prism 8.01 (GraphPad Software, San Diego, CA, USA). All data are presented as mean with standard deviation (SD).

## Results

### *S. rebaudiana* extracts antimicrobial studies

The tested plant extracts were characterized with a dry mass of: SRa – 32.8433 % (328.433 g/L) and SRm – 21.4921 % (214.921 g/L). In order to test the effects of the extracts on bacterial cells, a microdilution assay was performed. The mixtures of plant extracts and bacterial cultures cultivated in static conditions resulted in varied effects, that were strictly dose-dependent (Fig. 1). The SRa extract resulted in a decreased bacterial biomass of *E. coli* DSM 613 and of *E. coli* DSM 5695, with the exception of three concentrations (6.25 % and 3.125 % stimulated bacterial counts, 1.56 % kept bacterial biomass at the control level – 0 %). SRa extract addition caused a decrease in *P. syringae* DSM 21482 cells density at 0.19 %, whereas for most concentrations the biomass was kept at control level (50 %, 12.5 %, 6.25 %, 3.125 %, 1.56 %, 0.78 %, 0.097 %, 0.024 %) or was stimulated (25 %, 0.39 %, 0.048 %, 0.012 %, 0.006 %, 0.003 %) (Fig. 1A). SRm extract also showed inhibitory effect on *E. coli* DSM 613 cells in the majority of the tested concentrations, except for concentrations at which SRm did not influence bacterial growth (6.25 %, 3.125 %, 0.024 %, 0.012 %, 0.006 %, 0.003 %). A similar effect was observed for the SRm extract and *E. Coli* DSM 5695 where the majority of the tested concentrations decreased the bacterial biomass, except for two stimulating concentrations (12.5 %, 6.25 %) and one neutral concentration (3.125 %) that did not alter bacteria multiplication. The results were very similar to those obtained with the SRa extract. The addition of the SRm extract to *P. syringae* DSM 21482 cells led to the extract having no effect on the cells biomass, with the exception of four concentrations – one inhibiting (3.125 %) and three stimulating *P. syringae* DSM 21482 (0.012 %, 0.006 %, 0.003 %) (Fig. 1B).

**Fig. 1.**
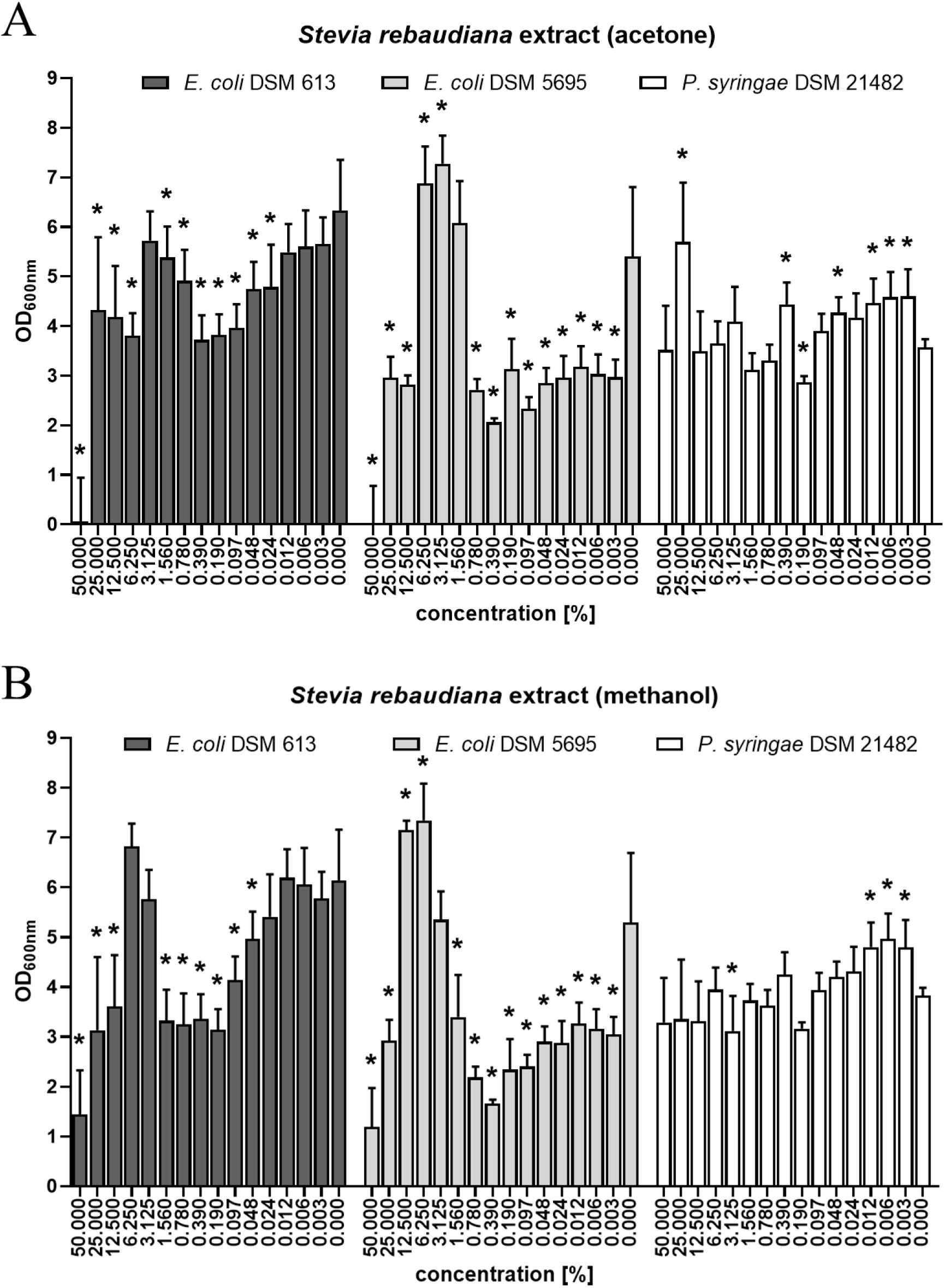
Activity of the *Stevia rebaudiana* extracts on bacterial cells expressed in OD changes of bacterial biomass. *Stevia rebaudiana* acetone extract activity assessed by microdilution method (**A**) and *Stevia rebaudiana* methanol extract activity assessed by microdilution method (**B**). Error bars represent standard deviation (SD) between samples. Means sharing the star asterisk are significantly different from the control (extract concentration 0 %) at p ≤ 0.05.

### Coincubation assay

In order to establish the influence of the plant extracts on phages activity and infection ability, a coincubation test was performed. The test was conducted in a static environment, in order to detect basic interactions between the tested factors. Coincubation showed varied results (Fig. 2). The addition of SRa extract did not significantly influence MS2 phage counts, with an exception of three concentrations (50 %, 25 %, 12.5 %) which resulted in a phage stimulating effect. When the SRa was added to the Phi6 phage lysate, the extract caused a phagicidal effect where no phage plaques were present in all tested concentrations. The combination of SRa extract with T4 phage resulted in mixed effects. Four SRa concentrations did not alter phage counts significantly (25 %, 3.125 %, 1.56 %, 0.097 %), two decreased phage PFU/mL (50 %, 6.25 %) and five stimulated T4 counts (12.5 %, 0.78 %, 0.39 %, 0.19 %, 0.049%) (Fig. 2A). Similarly to the SRa extract, SRm extract did not significantly influence MS2 phage counts, with the exception of two concentrations (50 % and 25 %) which stimulated the number of phage plaques. Also, the addition of the SRm extract to the Phi6 phage caused a significant phagecidal effect for all of the tested concentrations, which was also the case for the SRa extract. When the SRm extract was incubated with T4 phage, only the lowest concentration (0.049 %) did not significantly alter the phage counts, whereas all higher concentrations caused a stimulating effect (Fig. 2B).

**Fig. 2.**
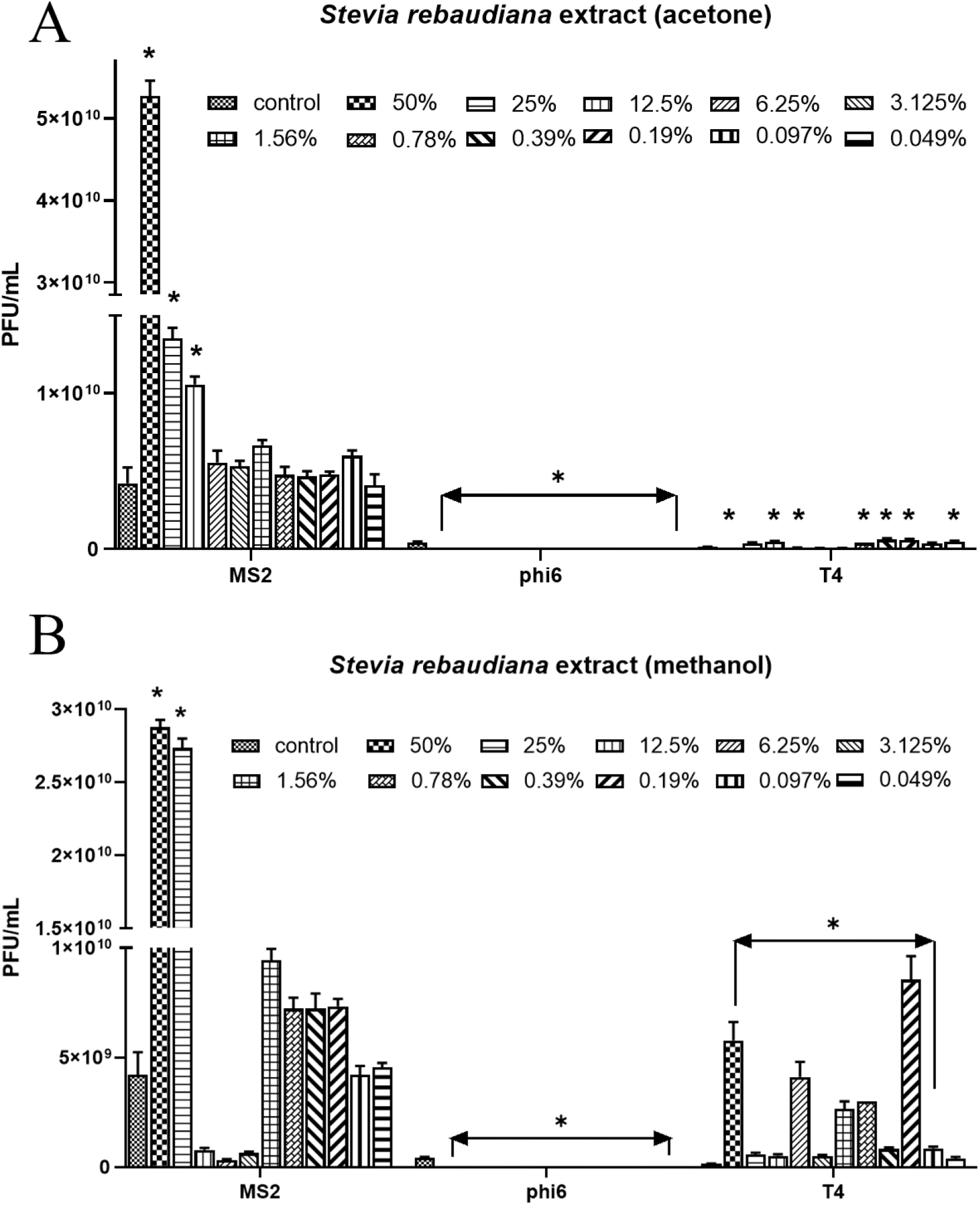
Phage titres (MS2, Phi6 and T4) after 24 h exposure to different concentrations of SR extracts. Phage counts after coincubation of phages with *Stevia rebaudiana* acetone extract (**A**) and after coincubation of phages with *Stevia rebaudiana* methanol extract (**B**). Error bars represent standard deviation (SD) between samples. Means sharing the star asterisk are significantly different from the control at p ≤ 0.05.

### Stoichiometries of the phage-extract interactions

For most of the tested samples a darkening of the sample was present, due to the dark colour of the concentrated extract. This was taken into account when analysing the results – the extracts optical density background was removed from the data for clear results interpretation.

The synograms experiment revealed complex relationships between the tested extracts and bacteriophages on their bacterial hosts (Fig. 3). Interestingly, in most of the cases the highest tested extracts concentration (25 %) caused increased growth of bacterial biomass, however after 24 h of incubation in the stationary environment the cells were no longer active (Fig. 3A-A’ – F-F’). This may indicate increased cell proliferation in the initial growth stage and earlier achievement of the stationary phase due to the presence of the extracts.

**Fig. 3.**
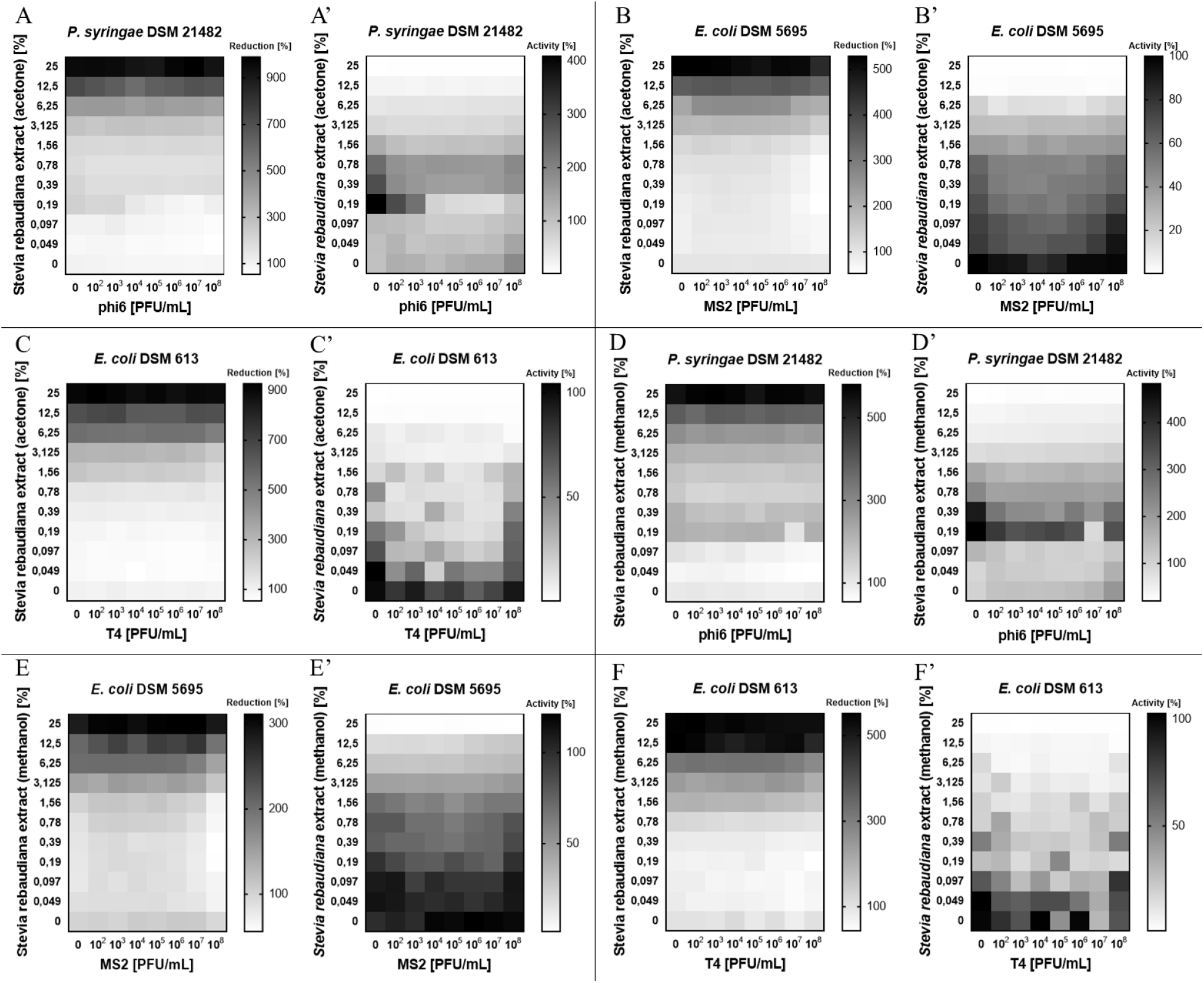
Results of phage-extract combinations on host bacteria. Paired heat-maps represent OD600nm measurements (**A**-**F**) and corresponding fluorescence measurements (**A’**-**F’**). Combined treatment of *Stevia rebaudiana* acetone extract with Phi6 phage (**A**-**A’**), MS2 phage (**B**-**B’**) and T4 phage (**C**-**C’**). Combined treatment of *Stevia rebaudiana* methanol extract with Phi6 phage (**D**-**D’**), MS2 phage (**E**-**E’**) and T4 phage (**F**-**F’**). Synograms (t = 24 h) represent the mean reduction (% of the control) or activity (% of the control) percentage of each treatment from three replicates.

Specific interactions between the tested elements were also observed. Three concentrations of the SRa extract (0.78 %, 0.39 %, 0.19 %) caused increased *P. syringae* cells activity, even though the OD values of the bacterial biomass were found to be dropping analogously to the decreasing concentration of the extract. Peak cell activity was detected in a mixture of 0.19 % SRa and no phage. Interestingly, at a SRa concentration of 0.049 % and maximum dose of phage (Phi6 – 10^8^ PFU/mL), the highest biomass reduction was detected, while cells activity grew by 44 % than compared to the control (Fig. 3A-A’). In general, the SRa extract influenced *E. coli* DSM 5695 by decreasing biomass proliferation (OD drops) analogously to the decreasing concentration of the extract, while at the same time cells activity rose almost to control levels. Here, an interesting phenomenon was also detected, when the highest biomass reduction was discovered (SRa: 0.19 % and MS2: 10^8^ PFU/mL), though cells activity level remained at 70 % compared to the control (Fig. 3B-B’). SRa extract combined with phage showed a similar influence on *E. coli* DSM 613 cells as on *E. coli* DSM 5695, however more scattered phenomena regarding cells activity were detected. To summarise, bacterial activity in some cases was extract-concentration and phage-concentration independent, and this mainly applied to peaks in cells activity, compared to the neighbouring samples. However, in one case (SRa: 0.049 % and T4: 10^4^ PFU/mL) the highest biomass reduction was discovered, along with the cells activity level showing a notably decrease: 20 % compared to the control (Fig. 3C-C’).

When SRm extract was used on *P. syringae* cells, the results obtained were very similar to those whern the SRa extract was added, four concentrations of the SRm extract (1.56 %, 0.78 %, 0.39 %, 0.19 %) increased bacterial cells activity, with simultaneous OD values of the biomass dropping analogously to the decreasing concentration of the extract. Peak cell activity was also detected in the mixture of 0.19 % SRm and no phage. Likewise, at a SRm concentration of 0.049 % and maximum dose of phage, the highest biomass reduction was detected, while cells activity grew by 62 % compared to the control (Fig. 3D-D’). A similar trend was also maintained for the results of the SRm extract and *E. coli* DSM 5695, compared to the use of the SRa extract under the same conditions. However, several cases of increased cells activity (compared to the control) were present. Interesting phenomenon (similar to that of the SRa experiment) was also detected when SRm extract was added: the highest biomass reduction was discovered at a SRm concentration of 0.19 % and MS2 phage concentration of 10^8^ PFU/mL, while the detected cells activity remained at 98 % compared to the control (Fig. 3E-E’). Similar tendencies were also present regarding *E. coli* DSM 613 and SRm extract, compared to the SRa extract – scattered phenomena in the case of cells activity were also detected. However, the most interesting phenomenon observed concerned SRm at 0.019 % and T4 at 10^8^ PFU/mL, resulting in the highest biomass reduction, along with a decrease of cells activity level, which was 16 % compared to the control (Fig. 3F-F’).

### The lytic performance of phages in the presence of plant extracts

To test the course and lysis ability of the phages in real time, phage performance against bacteria was tested in the presence of the extracts at different concentrations, measured by changes in optical density of the samples that were cultivated in the bioreactors with mixing (150 rpm) (Fig. 4). The extracts concentrations were chosen on the basis of results gained from the synograms experiment (combinations in which the described phenomena were present with maximal treatment doses as controls). Experiment time was also shortened (16 h), as the synograms experiment (>18 h) revealed unactive cells. For most of the tested samples, the darkening of the sample was present due to the dark colour of the concentrated extract. This was taken into account when analysing the data – the extracts optical density background was removed from the curves for clear results interpretation.

**Fig. 4.**
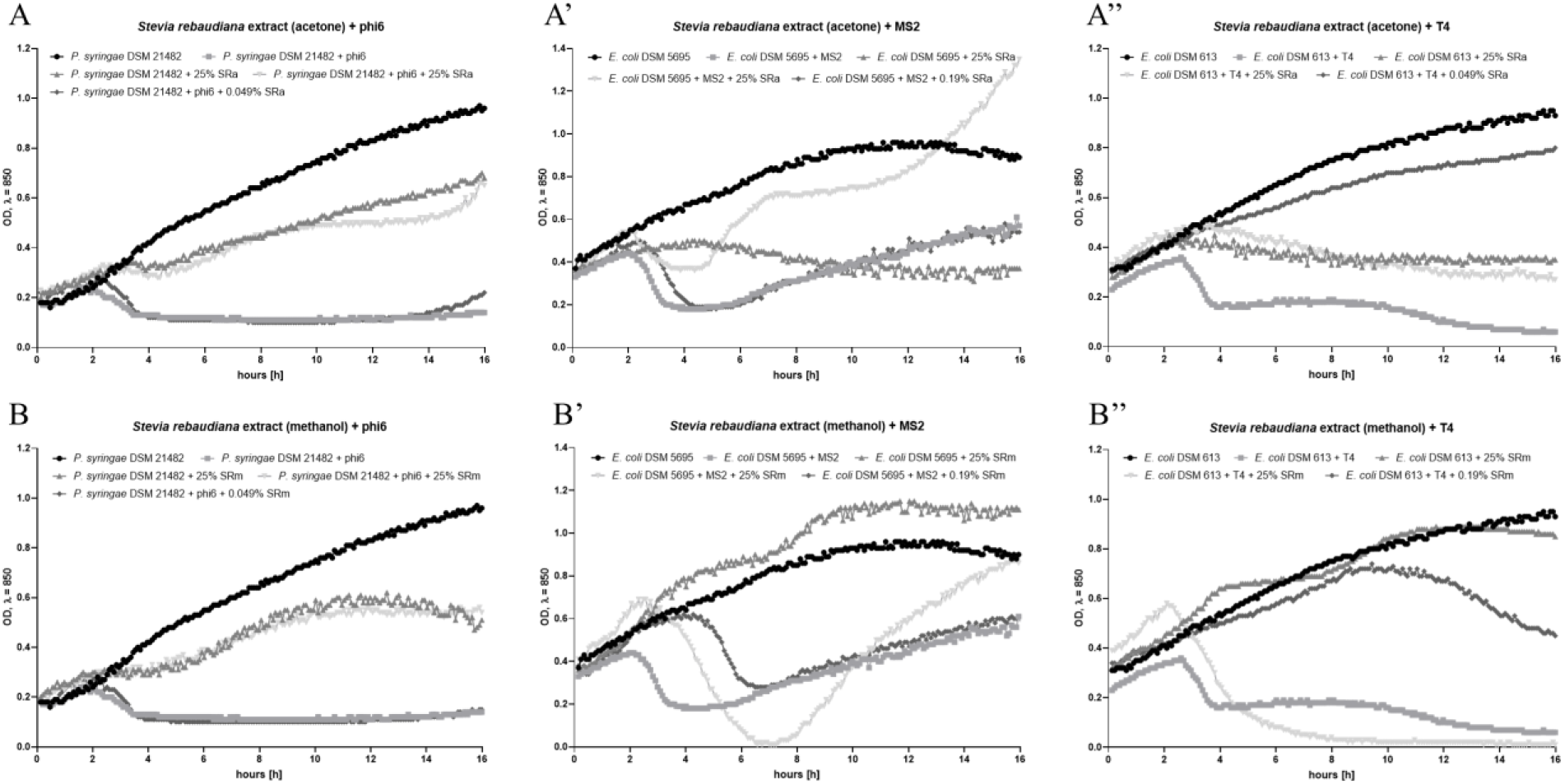
Bacteria growth curves treated by lytic phages and different extracts concentrations chosen in the synograms assay. Combined treatment of phages with of *Stevia rebaudiana* acetone extract (Phi6 - **A**; MS2 - **A’**; T4 - **A’’**) and combined treatment of phages with *Stevia rebaudiana* methanol extract (Phi6 - **B**; MS2 - **B’**; T4 - **B’’**).

The tested extracts influenced bacterial cells and phage activity in a varied manner, however, some dependencies were noted (Fig. 4). In general, high concentrations (25 %) of the tested extracts resulted in decreased phage lytic efficiency in most cases, either through a stimulating effect on bacterial cells cancelling the lytic effect of the phages or by influencing the phages alone, deactivating phage particles. The combination of *P. syringae* and SRa at a concentration of 25 %, with or without phage Phi6, resulted in a decrease in OD, lowering the growth curves – phage addition did not decrease the OD further. In addition, the curves were still in a rising stage, therefore bacterial cells were still multiplying. However, when the SRa concentration was 0.049 %, this led to the Phi6 and control lysis curve to be the same (Fig. 4A). The mixture of *E. coli* DSM 5695 and 25 % SRa extract caused a clear, gradual decrease in OD measurements, showing the extract’s toxic influence on bacteria. On the other hand, introducing a MS2 phage into this mixture resulted in a lytic effect up to5 h of the incubation. Afterwards, bacterial growth rebounded, growing rapidly. The reduction in SRa concentration to 0.19 %, led to the bacterial lysis curve almost identical to the control lysis curve, however lysis began a little later (after 2.5 h vs. controls 2 h) (Fig. 4A’). When SRa 25 % was added to the *E. coli* DSM 613 cells, this resulted in a visible decrease in OD, lowering growth curves, regardless of whether the T4 phage was used or not – which was similar in the case of *P. syringae*. However, in this case the curves were in the gradual reduction stage. When a SRa concentration of 0.049 % was used, it led to a slight reduction in the cell growth curve, compared to the growth control curve (Fig. 4A’’). The application of SRm extract at 25 % to the *P. syringae* suspension, offered almost the same result as when SRa extract was used. Regardless of the presence of the Phi6 phage, both growth curves were clearly lowered. However, after 12 h of incubation, the OD values started to fall gradually. When the SRm concentration of 0.049 % was applied, it also led to the Phi6 phage lysis curve to be the same as a control lysis curve, as when SRa was the tested extract (Fig. 4B). 25 % of SRm extract addition to the *E. coli* DSM 5695, caused a visible increased proliferation of the bacteria, stimulating the cells. When MS2 phage was added into this mixture, it led to a very strong lytic effect from 2.5 h to 7 h of incubation (a stronger and longer lasting effect than lysis control). However, later a strong bacterial proliferation was observed. Lowering the SRm concentration to 0.19 %, resulted in delayed but elongated phage lysis, compared to the control curve (control from 2 h till 3.5 h = 1.5 h; tested sample from 4.5 h till 6.5 h = 2 h) (Fig. 4B’). The addition of SRm at the concentration of 25 % to the *E. coli* DSM 613 cells did not generate any specific effect on the bacteria – the growth curve was almost identical to the bacterial control growth curve. When bacteriophage T4 was also added, this was similar to the case of MS2 phage and *E. coli* DSM 5695, a marked strong lytic effect was observed. The effect was also longer and stronger than control – lytic activity was detected from 2 h of incubation, and no bacterial growth rebound was observed. Lowering SRm concentration to 0.19 % caused the curve to be highly comparable to the control growth curve, until 9 h of incubation. After that time, the curve began to drop, indicating the death of the bacterial cells (Fig. 4B’’).

## Discussion

Stevia is mainly known as a natural sweetener, it is also gaining interest due to the many beneficial effects not only for humans (25, 32, 11), but also animals, where these herbs can serve as feed additives (having a supplementary role) or alternative feed source (11, 4, 5, 27). It has also been shown, that phages can be used as feed additives to reduce bacteria in animal preslaughter, without negatively impacting microbial communities (40). Mindful of this, we tested possible interactions between *Stevia rebaudiana* acetone and methanol extracts, and lytic phages as antibacterial agents in an environment of bacteria related to fauna and flora diseases.

SRa and SRm extracts revealed diverse and multidimensional interactions with the mixtures of phages and their bacterial hosts. In order to explain these interactions, it is necessary to draw general conclusions from the experiments performed and later propose hypotheses explaining the results.

The lysis profile experiment of *P. syringae* and SRa at a concentration of 25 %, with or without phage Phi6, resulted in OD drops and when the SRa concentration was 0.049 %, this led to the Phi6 and control lysis curve to be the same. These concentrations stimulated the cells in the modified MIC test, but proved to be phagicidal (tests in static conditions – no mixing). It is possible, that the introduction of dynamic conditions (mixing) neutralised extracts stimulating influence on bacteria, and the phagicidal effect in a concentration-dependent manner. When *E. coli* DSM 5695 was tested, 25 % of the SRa extract caused a gradual decrease in OD, while after a MS2 phage was added, the lytic effect was observed up to 5 h of the incubation followed by bacterial growth rebound. SRa extract at 0.19 %, led to the lysis and control curve to be almost identical. These concentrations were toxic to cells in the MIC test, and SRa 25 % stimulated MS2 phage. In this case, we also hypothesise that dynamic conditions neutralised the extracts’ influence on the bacteria and phage-inducing effect, though to a lesser extent. When SRa 25 % was added to the *E. coli* DSM 613 cells, it resulted in a similar outcome as in the case of *P. syringae* – a visible decrease in OD was noted, regardless if the T4 phage was used or not. Also, a SRa concentration of 0.049 % led to only a slight reduction in the cell growth curve, compared to the growth control curve. These concentrations were toxic to cells in the modified MIC test, and SRa 0.049 % stimulated the phage. Here, we also would argue that to a lesser extent, dynamic conditions neutralised the extracts influence on bacteria in a concentration-dependent manner. Moreover, the phage-inducing effect was not visible due to the small amount of phage particles (10^4^ PFU/mL). The application of 25 % of the SRm extract on *P. syringae* cells gave almost the same result as when a SRa extract was used – regardless of the presence of the Phi6 phage, both growth curves were clearly lowered. SRm at the concentration of 0.049 % with the addition of Phi6 phage caused the lysis curve to be identical as the control lysis curve, the effect was the same when SRa was used. These concentrations did not affect cells in the modified MIC test, but were phagicidal. In this case, dynamic conditions could also be the answer for those results. It caused the extract to be slightly cell-growth inhibiting at 25 %, and also in a concentration-dependent manner neutralised the phagicidal effect. 25 % of SRm extract addition to the *E. coli* DSM 5695, caused increased bacterial proliferation. With the addition of MS2 phage, a lytic effect was observed that was overall stronger and longer than compared to lysis control. However, bacterial growth rebound was detected. Lowering SRm concentration to 0.19 %, resulted in delayed but also elongated phage lysis, compared to the control curve. These concentrations influenced the cells toxically in the modified MIC test, and 25 % SRm was phage stimulating. In addition, mixing caused the extract to lose its toxic influence on cells (the effect even shifted to cell-stimulating) but the effect on MS2 phage remained to be inducing at 25 % SRm. When *E. coli* DSM 613 was tested, the addition of 25 % SRm did not generate any specific effect on the bacteria, but when phage T4 was added, a stronger and longer lytic effect was noted than the lysis control which was similar to what occurred in the case of the MS2 phage and *E. coli* DSM 5695, but with no bacterial growth rebound. Lowering the SRm concentration to 0.19 % caused the curve to be highly similar to the control growth curve, but after 9 h it began to drop. These concentrations inhibited cell growth in the modified MIC test, and stimulated MS2 phage. Therefore, the results presented here are also a consequence of the extracts losing their toxic activity on *E. coli*, but the effect on T4 phage remained to be inducing in a concentration-dependent manner.

In general, the literature shows varied effects of stevia on bacterial cells, especially in the case of human microbiota – from inhibitory activity to stimulating effects (9, 38, 31). However, inconsistencies have also been found in research concerning standard laboratory bacterial strains, mainly caused by different extract types. *Stevia rebaudiana* water, methanol, ethyl acetate and hexane extracts that were examined against few selected microorganisms revealed, that the water extract was only active against *B. subtilis* and *S. aureus* and the methanol extract was the most active against *P. aeruginosa. E. coli* proved to be most susceptible to hexane extract, and subsequently to ethyl acetate and methanol extract, with no effect when water extract was used (37). Within other author’s research, where water, ethanol, petroleum ether, cyclohexane, acetone and chloroform extracts were tested, petroleum ether extract was found to inhibit the growth of *E. coli* completely. The highest antibacterial index was also obtained for petroleum ether extract against all pathogens, with the highest antimicrobial activity against *S. aureus, E. faecalis* and *P. aeruginosa. E. coli, P. mirabilis* and *B. subtilis* were the most susceptible to water, ethanol and acetone extracts (12). When stevia extracts were obtained through the Ayurvedic Pharmacopeial Method (water and alcohol), the Soxhlet method and column extraction, the highest rate of susceptibility was exhibited by *E. aerogenes* for all the extracts. Alcohol extract showed the highest activity against all tested bacteria. *E. coli* was visibly inhibited by alcohol and Soxhlet extracts, the column extract was active to a lesser extent and the water extract was not active at all (21). Other experiments revealed that acetone extract had most effective antibacterial potential, followed by ethyl acetate extract. The acetone extract showed greater activity against Gram-positive rather than against Gram-negative organisms. *E coli* was equally susceptible to ethyl acetate and acetone extracts, with no influence noted by water extract (16). Contrary to those results, it was also shown that methanolic extract was the most effective against all the tested bacteria. *E. coli* susceptibility to chloroform and methanol extracts was similar, and no effect was observed when the water extract was applied. However, it is worth noting that it was also established, that chloroform and methanol extract exhibited concentration dependent antibacterial inhibition. Furthermore, it was also observed that when extracts were diluted, inhibition was greater in some cases (22). Difficulty in the interpretation of results and comparison is even greater when another variable is taken into consideration. It has been shown, that even the drying methods of *Stevia rebaudiana* can affect its quality and antimicrobial activity, leading to longer lasting bacteria inhibition (19). Unfortunately, all of the presented data were collected by simple diffusion methods, therefore it is also possible that the results obtained by other authors could present different outcome when using microdilution or real-time growth methods. Even if there are no papers describing the influence of stevia on bacteriophages, the antiviral activity of the plant was confirmed – hot water extracts of stevia showed anti-human rotavirus (HRV) activity. Stevia inhibited the replication of all four serotypes of HRV *in vitro*. It was indicated that the inhibitory mechanism is a blockade of virus binding and inhibitory components were heterogeneous anionic polysaccharides with different ion charges (9). These findings support our research with regards to SRa and SRm activity, as well as a dynamic environment thesis – charged particles could behave differently in mixing conditions rather than in static.

Currently, the is limited information in literature regarding the effects of extracts on phages or simultaneous application of both. Even fewer works describe these interactions in the environment of bacterial hosts. Previous scientific works have mainly described the plaque-forming ability of the phages after their contact with plant extracts (6-8, 17). However, there are some findings that describe more complex interactions. One of these shows the influence of *Phoenix dactylifera* L. acetone extract on lytic *Pseudomonas* phage – the extract inhibited phage infectivity and completely prevented bacterial lysis, but no information about extracts influence on bacteria was presented (15). Some plant extracts have also shown concentration-dependent antiviral activity and a reduction of phage yield (2). The importance of performing multiple tests in order to understand such complex interactions that can change in different environments, was noted in another study. It was found that black cumin extract increased phage plaque size, however this effect was not reflected in phage titers in a liquid medium. In general, in liquid media experiments, no synergistic effects were detected. Furthermore, the observed interactions were more closely related to antagonism, something we also noted in our work (39). Moreover, plant extracts and lytic phages can significantly reduce bacterial concentrations compared to untreated and extract treated controls, but these reductions do not extend further (decrease over time) (26).

Through our work, we hypothesise that changing environment conditions by mixing can be responsible for some of the observed effects of the stevia extracts. It is worth pointing out that in our previous article we also observed plant extracts stimulation of bacteria, after testing the mixtures in a dynamic environment of a bioreactor (24). This phenomenon could be explained by the enhanced physical contact of the mixtures molecules due to mixing, which in consequence enhances the interactions therein, not observed in static conditions. However, a thorough understanding of this phenomenon may require further research.

## Conclusions

The effects of *S. rebaudiana* acetone (SRa) and methanol (SRm) extracts on the course of phage lysis and activity in the dynamic environment depend on the species of the phage and bacterial host. The greatest differences in the used extracts are noted for *E. coli* strains and their phages, whereas *P. syringae* and Phi6 phage reacted similarly to the SRa and SRm extracts. Differences also emerge for a single extract (SRa or SRm) within *E. coli* strains and their phages – therefore one should test each extract type on a case-by-case basis and not generalize.

Dynamic conditions can alter the influence of extracts on bacterial cells and phages, also in a concentration-dependent manner. Because the interactions of phage-extract factors against bacteria in a static environment are often different than in a dynamic environment, many varied experiments should be performed, especially when examining multifactorial mixtures. Further studies are needed to understand the basics of the interactions between phages and plant extracts for the possible future use of phage-extract combinations as antibacterial mixtures.

An appropriate concentration of the selected herb extract may have antibacterial properties. It may also increase the activity of phages. Due to the higher activity of phages containing the selected extract than simply the phages themselves, these mixtures could be used in multi-drug resistant bacterial infections treatments.

## Conflict of Interests Statement

The authors declare that there is no conflict of interests regarding the publication of this article.

## Financial Disclosure Statement

This research did not receive any external funding.

## Animal Rights Statement

Not applicable.

## Notes

### Competing Interest Statement

The authors have declared no competing interest.

